# In a nutshell, a reciprocal transplant experiment reveals local adaptation and fitness trade-offs in response to urban evolution in an acorn-dwelling ant

**DOI:** 10.1101/2020.08.21.251025

**Authors:** Ryan A. Martin, Lacy D. Chick, Matthew L. Garvin, Sarah E. Diamond

## Abstract

Urban-driven evolution is widely evident, but whether these changes confer fitness benefits and thus represent cases of adaptive urban evolution is less clear. We performed a multi-year field reciprocal transplant experiment of acorn-dwelling ants across urban and rural environments. Fitness trade-offs via survival were consistent with local adaptation: we found a survival advantage of the ‘home’ treatments compared to the ‘away’ treatments. Seasonal bias in survival was consistent with evolutionary patterns of gains and losses in thermal tolerance traits across the urbanization gradient, such that rural ants in the urban environment were more vulnerable in the summer, putatively due to low heat tolerance and urban ants in the rural environment were more vulnerable in winter, putatively due to an evolved loss of cold tolerance. The results for fitness via fecundity were more complex. Fecundity differences were present in the rural environment, but not the urban environment, and could reflect mismatched cues for the seasonal production of reproductives. To broadly contextualize our results, we performed a multi-species meta-analysis of urban adaptive evolution studies and found general support for local adaptation. The rural adaptation signal was stronger than for urban adaptation, consistent with the relative differences in time for adaptation to occur.

## Introduction

There is growing evidence that urbanization influences the evolution of species that inhabit and interact with urban environments (Johnson and Munshi-South 2017; Schilthuizen 2018; Szulkin et al. 2020b). Some of these evolutionary responses stem from changes in gene flow and genetic drift due to anthropogenic barriers and corridors (Miles et al. 2019). Others have been attributed to selection driven by changes in the physical (*e.g.*, substrate, temperature, flow regime; noise/light pollution, environmental toxins) and biotic (*e.g.*, population and species interactions) environment (reviewed in Johnson and Munshi-South 2017). With a few notable exceptions (*e.g.*, Whitehead et al. 2012; Reid et al. 2016), these latter examples of urban evolution generally provide strong, but indirect evidence that phenotypic divergence in cities is indeed adaptive. For example, Winchell et al. (2016) found that limb length and toepad lamellae structure had likely evolved between urban and rural *Anolis* lizards, and that these changes were correlated with locomotor performance (Winchell et al. 2018). However, locomotor performance has not been linked to fitness differences or measured under natural conditions in these populations. Other studies in the same *Anolis* populations (Campbell-Staton et al. 2020), and in white clover (Thompson et al. 2016) have shown allele frequency shifts between urban and rural populations, likely in response to urban heat island effects (also see Müller et al. 2013; Harris and Munshi□South 2017; Kerstes et al. 2019). However, the fitness effects of these changes have not yet been measured for these systems. The strength and direction of selection has also been measured across urbanization gradients (Lambrecht et al. 2016; Irwin et al. 2018; Start et al. 2018), but selection analyses, on their own, are not sufficient to demonstrate local adaptation. The fitness effects of increased thermal tolerance to urban heat islands have been measured for water fleas (Brans and De Meester 2018), damselflies (Tüzün et al. 2017), and acorn ants (Diamond et al. 2018a), as have the fitness consequences of dispersal in fragmented urban landscapes (Cheptou et al. 2008), but each of these examples took place in the laboratory or artificial conditions.

Cities provide an unplanned, experimental arena for better understanding the process of local adaptation (or their lack thereof) to novel and changing environments (Diamond and Martin 2020a). While it is now clear that rapid evolutionary responses, taking place within the timescale of human lifespans, routinely occur in natural populations (Hendry and Kinnison 1999; Reznick et al. 2019), we have much to learn about if or how such contemporary adaptation unfolds. For example, does adaptation to novel environments commonly impose fitness trade-offs in relation to the ancestral environment? Stronger fitness trade-offs between environments might facilitate population divergence and speciation, but they could also impede adaptation to novel environments (Kawecki and Ebert 2004; Hendry 2009). And how (mal)adapted are populations in novel or changing environments in comparison to related populations inhabiting ancestral conditions? Such research questions are especially relevant in light of the current pace, magnitude, and ubiquity of anthropogenic change (Brady et al. 2019; Diamond and Martin 2020a).

By comparing the relative fitness of populations in their native environment versus a foreign environment, and by comparing a native population’s fitness in their home environment to that of a foreign population, reciprocal transplant experiments are a powerful approach for measuring local adaptation and identifying potential fitness trade-offs (Kawecki and Ebert 2004; Blanquart et al. 2013). While this fact is well-recognized in the emerging field of urban evolutionary biology (Donihue and Lambert 2015; Rivkin et al. 2019), reciprocal transplants between urbanized and non-urbanized habitats are still rare. Moreover, the urban-focused reciprocal transplants carried out to date provide negative evidence (*e.g.*, Capilla-Lasheras et al. 2017) or mixed evidence (*e.g.*, Gorton et al. 2018) for urban adaptation, leaving open the question of how common adaptive evolution to city environments might be.

In this study, we set out to address these gaps in our understanding of urban local adaptation. Our study includes two components: a multi-year reciprocal transplant experiment using an acorn-dwelling species of ant, *Temnothorax curvispinosus*, and a meta-analysis of urban local adaptation across multiple taxa. Previous multi-generation common garden experiments with acorn ants have demonstrated evolutionary divergence in thermal tolerance between urban and rural populations in response to urban heat island effects (Diamond et al. 2017, 2018a; Martin et al. 2019). These shifts were suggested to reflect local adaptation on the basis of selection (via fecundity) for greater heat tolerance and fitness trade-offs between urban and rural populations across urban versus rural-mimicking temperature treatments in the laboratory (Diamond et al. 2018a).

The reciprocal transplant experiment allowed us to robustly test for fitness trade-offs and evidence of local adaptation. We transplanted urban and rural acorn ant colonies across multiple urban and rural sites in Cleveland, Ohio, USA, and monitored colonies for over two years to assess colony survival and fecundity. Note that for acorn ants, ‘rural’ sites refer to typical forested sites for this species, whereas ‘urban’ sites refer to forest islands embedded within the urban matrix (see Diamond et al. 2018b); hereafter, we refer to our sites as urban versus rural for simplicity and consistency with other research in this area. We also phenotyped a subset of colonies for heat and cold tolerance traits. If local adaptation is present, we expected transplanted urban ants to have the highest fitness in the urban environment and rural ants to have highest fitness in the rural environment. Further, if local adaptation leads to fitness trade-offs across environments, we expected that populations will have lower fitness in their non-native environments than in their native environments. Because we were able to census colony survival and fecundity over multiple time points, we also examined the evidence for environmental temperature as a selective agent across winter and summer seasons in urban and rural environments. Finally, using these same approaches and inferences, we broadened our analysis of urban local adaptation with a meta-analysis. Owing to the scarcity of urban field reciprocal transplant studies, we considered both field transplant studies and laboratory experiments. Using this broader range of taxa, including both plant and animal study systems, we again examined the evidence for local adaptation by assessing whether fitness was highest in the ‘home’ environment.

## Materials and methods

### Study system

We used the small, acorn-dwelling ant, *Temnothorax curvispinosus*, to explore the potential for local adaptation to an urban heat island. The entire colony including the queen, workers, and brood (immature ants in the egg, larval and pupal stages of development) resides within the acorn environment. There can be multiple queens, especially as colonies mature, often driven by daughters of the reproductive caste remaining in the nest rather than dispersing (Stuart 1987). The queen(s), brood, and young workers that are typically tasked with brood care, remain entirely within the nest environment. Older workers are typically the only ants that regularly leave to gather food resources and return these items to the nest. Because of the unique nesting structure of acorn ants, they are highly amenable to being transplanted between different sites. Further, the colony remains aboveground in the acorn environment for the entire year, which not only facilitates collection of whole colonies, but also ensures that acorn ants experience near-surface temperatures. Acorn ants do not have the capability to retreat underground like many ant species (Herbers 1989; Herbers and Johnson 2007; Mitrus 2013), and are thus unable to use this mechanism to avoid stressfully warm temperatures associated with the urban heat island effect or stressfully cool temperatures associated with winter conditions.

The lifespan properties of components of the acorn ant colony also make it highly conducive to long-term reciprocal transplant studies, which are the most direct method to assess local adaptation. As a colony unit, acorn ants are remarkably long-lived, owing to the fact that queens can live five or more years (Keller 1998; Negroni et al. 2019). By contrast, worker lifespan is considerably shorter, on the order of one or more months (Modlmeier et al. 2013; Penick et al. 2017). Queens must continually produce new eggs to replace old workers in nest-tending and foraging tasks, as the queen does not forage herself. The production of new workers is also important for achieving a large enough colony size that investment shifts from generating non-reproductive workers (colony maintenance) to reproductive alates, or winged, sexual, male and female ants.

### Reciprocal transplant experiment

The reciprocal transplant experiment was conducted in urban and rural tree-covered sites of Cleveland, Ohio, USA (see **Supplementary Material** for details of transplant pen construction). We sourced acorn ants from four urban sites and two rural sites (**Table S1**). We used percent developed impervious surface area (ISA) from the National Land Cover Database (Yang et al. 2018) as the criterion for designating urban and rural sites. Urban sites were defined as those with ISA values between 40 and 61% and rural sites were defined as those with ISA values of 0%. Mean ISA values for each site were calculated with a 120 m buffer using the focal statistics tool in the spatial analyst toolbox in ArcGIS 10.4. Urban and rural acorn ant microclimates differ by more than 4 °C during the main activity season for acorn ants (Diamond et al. 2018a). Colonies were collected from field sites between 22 May and 19 July 2017. We focused colony collections relatively early in the year to avoid inadvertently splitting up polydomous single colonies. Acorn ant colonies typically overwinter in a single acorn and, for mature colonies, typically expand to multiple acorns during the later summer months (Alloway et al. 1982).

All colonies were held in a laboratory growth chamber (Panasonic MIR 154) at a constant 25 °C (14:10 L:D) for a minimum of three days. During this period, colonies were provided with unlimited access to water, sugar solution (25% sucrose), and dead mealworms. We counted the number of queens, workers and brood in each colony. At the end of the laboratory acclimation period, we assessed heat and cold tolerance traits of worker ants from a subset of the colonies that had sufficiently large colony sizes, *i.e.* for which the thermal tolerance trials would not eliminate the current worker pool. Colonies that underwent thermal tolerance testing had to contain at least 20 workers that were transplanted into the field with the queen and brood meaning colonies had to contain at least 30 workers upon initial collection to allow 10 to be tested for thermal tolerance traits. The thermal tolerance data allowed us to examine patterns of phenotypic selection on thermal tolerance traits via colony survival to the end of the experiment. We assessed five individual workers for heat tolerance and a different five individuals for cold tolerance per colony. Owing to logistical constraints, we were able to do this for 12 urban and 9 rural colonies, representing 48% of the total number of initial colonies in the experiment. Two workers from a single rural colony were injured during the assessment of heat tolerance, and so these data points were excluded from the experiment, leaving a total of 118 individual estimates of heat tolerance and 120 estimates of cold tolerance. Our measures of heat and cold tolerance were the critical thermal maximum (CT_max_) and critical thermal minimum (CT_min_). These assays were performed using a dynamic temperature ramping protocol, raising or lowering the temperature by 1 °C min^−1^ until the loss of muscular coordination. Starting temperatures were 34 and 16 °C for heat and cold tolerance, respectively. Note that the thermal tolerance assays were destructive, preventing the return of these worker subsets to the colony, but ten workers represents only 13% of the mean colony size in our experiment and is thus highly unlikely to undermine colony performance. Because our colony collection dates spanned late May to early July, we examined the potential for seasonal acclimation effects on thermal tolerances of urban and rural population acorn ants. We found no evidence for such effects (**Table S2**) and thus did not further consider the influence of the timing of collection on thermal tolerance.

Acorn ant colonies were transplanted into four urban and two rural sites between 25 May and 31 July 2017 (**Table S1**). Although the two rural transplant sites were the same as the rural colony collection sites, two of the four urban transplant sites differed from the colony collection sites. This difference simply reflected the confluence of variation in colony density at different urban sites and our ability to obtain permissions for performing reciprocal transplants in urbanized habitats. Colonies were transplanted to home treatments (urban source to urban habitat and rural source to rural habitat) or away treatments (urban source to rural habitat and rural source to urban habitat). For the treatment groups rural-origin ants into rural environments and urban-origin ants into rural environments, each group had 10 colonies initially. For the treatment groups rural-origin ants into urban environments and urban-origin ants into urban environments, each group had 12 colonies initially. For the home treatment transplants, colonies were always transplanted into a new site of the same habitat designation rather than the same site of origin. Collection and transplant sites were each already known to already harbor acorn ant colonies to ensure appropriate microhabitat conditions within the experiment. A given colony was transplanted intact as a single colony unit including the original queen, workers and brood that were in the colony upon the initial collection from the field site, *i.e.*, we did not split colony members across treatments or sites. Only one colony was placed within each individual transplant pen to allow tracking of colony survival and size on a per-colony basis. Initial colony size with respect to the total number of queens, workers, and brood were comparable across the urban and rural source habitats prior to transplantation (**Table S3**).

The experiment began on 25 May 2017 with the first transplantations of colonies to their field treatments and ended on 12 August 2019 with the recovery of the surviving colonies from the transplant pens. Over this period, colonies were censused 11 times (**Fig. 1**). At these census points, each pen was assayed for whether the colony had survived. Surviving colonies were briefly brought from the field into the laboratory to be assessed for colony size including the number of queens, workers and brood. Reproductive effort (fecundity) was also assessed by counting the number of alates produced. If alates were found, they were removed from the colony prior to returning the colony to the field. We did this to prevent genetic contamination by dispersants from the ‘away’ transplants. Colonies were allowed to regather their nest inside their acorn before being returned to their transplant pen. All colonies were returned to their original pens in which they began the experiment.

**Figure 1.**
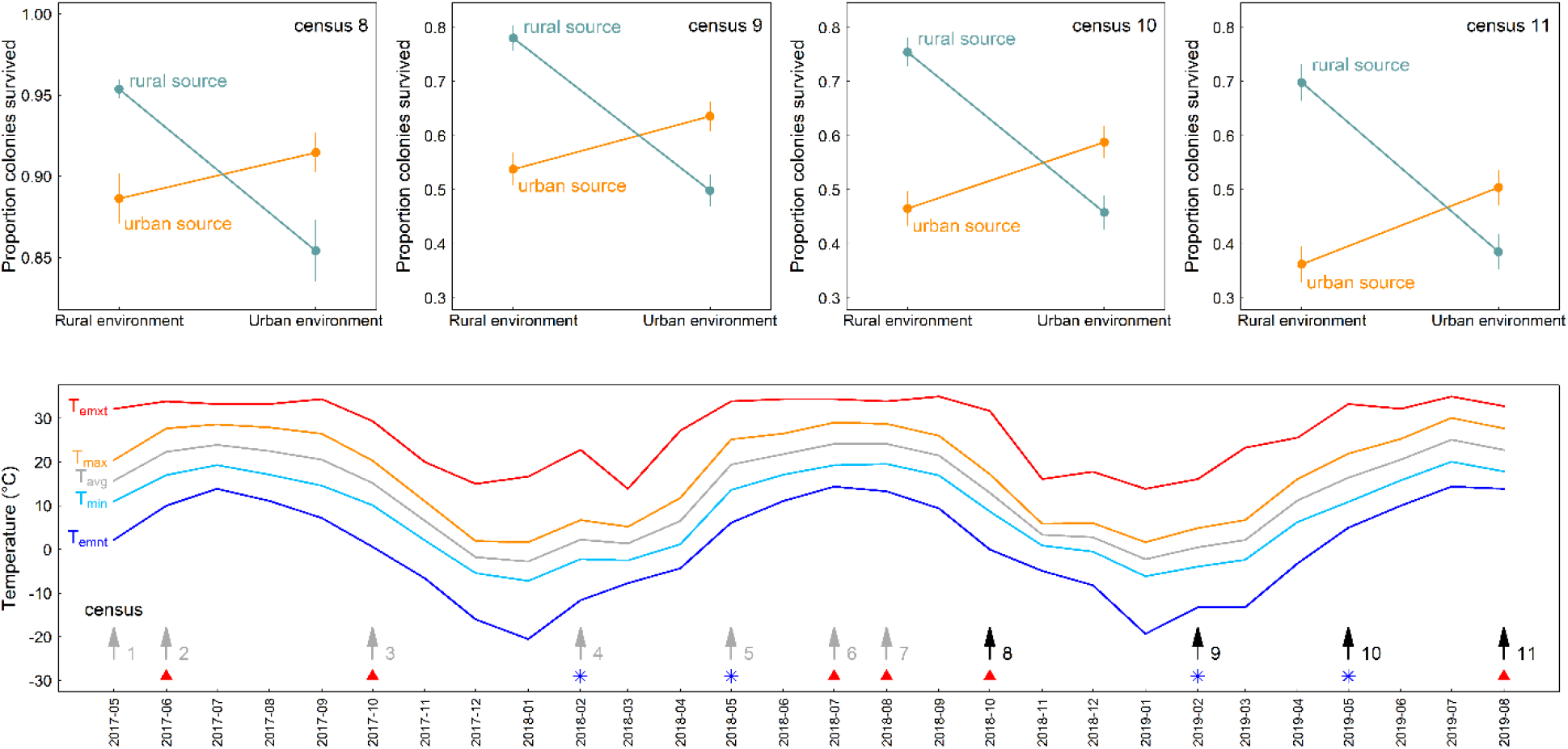
Fitness tradeoffs for the final four census points (census 8 through 11) (top row), and the monthly temperature profile with all census points (census 1 through 11) indicated (bottom row). Fitness tradeoff plots include estimated mean survival ± 1 SE for each combination of source population and environment from a generalized linear model that accounts for temporal autocorrelation. Note that census 8 y-axis is scaled from 0.825 to 1, whereas censuses 9-11 are all scaled from 0.3 to 0.825 owing to high overwinter mortality following census 8. On the temperature profile plot, arrows indicate the timing of each census (at the monthly resolution); black arrows correspond with the census points highlighted in the top row. Red triangles indicate a warm-season census (predominantly summer into fall) and blue asterisks indicate a cool-season census (predominantly winter into spring), that correspond with the designations used in the analysis of seasonal bias in between-census survival (**Fig. 3**). These symbols refer to the period between the previous and focal census point, *e.g.* the red triangle at census 3 refers to a warm period between census 2 and 3 whereas the blue asterisk at census 4 refers to a cool period between census 3 and 4. Temperature indices include average monthly temperature (T_avg_), the extreme monthly minimum and maximum temperatures (T_emnt_ and T_emxt_), and the average monthly minimum and maximum temperatures (T_min_ and T_max_). Dates are reported as year-month.

The multi-year and multi-census aspects of our reciprocal transplant experiment allowed us to explore not only the experiment endpoint, *i.e.* fitness differences among warm urban source population ant colonies vs. cool rural source population ant colonies across environments, but also the seasonal impacts of climate on ant colony survival and colony size. Because we were interested in broad-scale patterns of ant responses to seasonal variation in climate, we used local weather station data at the monthly resolution (GSOM – NOAA; station ID: GHCND:USW00014820) which best matched our colony censusing frequency. Our focal temperature variables included the monthly overall average, average maximum, average minimum, extreme maximum and extreme minimum temperature. All climatological data were extracted using the *ncdc* function from the {rnoaa} package (Chamberlain 2019).

### Meta-analysis of local adaptation

To find reciprocal transplant experiments measuring fitness in urban versus rural habitats we searched through the database of urban evolution studies compiled by Johnson and Munshi-South (2017) and searched ISI Web of Science using the keywords (1) ‘reciprocal transplant’ and ‘urban evolution’; and (2) ‘fitness’ and ‘urban evolution’. Note that we use ‘urban’ versus ‘rural’ in a general sense. Specific site characteristics must be interpreted within the specific study system. As such, we relied on the study authors’ definitions of what constituted an urban or rural (alternatively, ‘non-urban’) site. We retained studies that (1) measured fitness as survival, fecundity or a composite measure of the two (*i.e*., total fitness); (2) reciprocally transplanted individuals between urban and rural habitats or exposed individuals from urban and rural populations to both urban and rural conditions (*e.g.*, temperature) in the lab. For lab studies with multiple treatments (*e.g*., more than two temperature treatments), we chose fitness measured in the treatments that best matched the natural urban and rural conditions for the study, as reported by the study authors and associated studies (**Table 1**). We did not retain studies that included additional experimental manipulations or that performed lab selection experiments. Using these criteria, we retained seven studies from which we extracted fitness measures and standard errors **(Table 1**). Standard errors were not available for two studies measuring survival (Kettlewell 1955, 1956), and so we calculated binomial errors from the raw frequencies using the *binom.confint* function in the {binom} library (Dorai-Raj 2014).

**Table 1.** Summary of studies included in the meta-analysis of local adaptation to urban environments.

| SPECIES | TREATMENTS | STUDY TYPE | FITNESS METRIC | REFERENCE |
| --- | --- | --- | --- | --- |
| *Ambrosia artemisiifolia* | urban-rural transplants | field | total fitness | Gorton et al. 2018 |
| *Biston betularia* | urban-rural release-recapture | field | survival | Kettlewell 1955 |
| *Biston betularia* | urban-rural release-recapture | field | survival | Kettlewell 1956 |
| *Coenagrion puella* | temperature (16°C, 20°C) | lab | survival | Tüzün et al. 2017 |
| *Cyanistes caeruleus* | urban-rural transplants | field | survival | Capilla-Lasheras et al. 2017 |
| *Daphnia magna* | temperature (20°C, 24°C) | lab | total fitness | Brans and DeMeester 2018 |
| *Temnothorax curvispinosus* | temperature (21°C, 29°C) | lab | fecundity | Diamond et al. 2018a |
| *Temnothorax curvispinosus* | urban-rural transplants | field | survival (final census), fecundity (cumulative) | current study |

### Statistical analyses

#### Cumulative survival

To evaluate our main hypothesis of whether ‘home’ treatments (urban-origin ants transplanted into the urban environment and rural-origin ants transplanted into the rural environment) exhibited greater fitness (survival) than ‘away’ treatments (urban-origin ants transplanted into the rural environment and rural-origin ants transplanted into the urban environment), we constructed a generalized linear model. As the response, we included cumulative survival to each census point, *i.e.* the fraction of colonies within a given treatment that were alive at a given census relative to the starting number of colonies in that treatment. As predictors, we included treatment, a four-level factor describing each combination of source population and environment, a continuous covariate for the census point, and the interaction between these two variables. We used a beta error distribution and included an arima correlation structure to account for temporal non-independence among successive census points. We used the *glmmTMB* function from the {glmmTMB} library (Brooks et al. 2017) in R (R Core Team 2019) to construct this model.

To analyze this model, we performed analysis of variance to determine the statistical significance of the fixed effect predictors including main effects of treatment, census and their interaction. We then performed post-hoc analyses using the *emmeans* function from the {emmeans} library (Lenth 2019) to determine the statistical significance of the difference between urban and rural populations within each environment type (urban versus rural environment) (*sensu* Hereford 2009) as well as the difference between urban and rural environments for each population (urban-origin versus rural-origin population). These comparisons allowed us to evaluate census-specific differences in survival and identify the census points at which populations and environments diverged with respect to cumulative survival.

#### Between-census survival

To evaluate our secondary hypothesis that there will be seasonal bias in mortality based on the nature of the ‘away’ treatment, *i.e.* that urban-origin ants transplanted to the rural environment will exhibit greater mortality in winter owing to the evolved loss in cold tolerance and that rural-origin ants transplanted to the urban environment will exhibit greater mortality in summer owing to the lack of evolved gain in heat tolerance, we used a general linear model. For this analysis, we computed the between-census survival for each treatment group, *i.e.* given the starting number of colonies at one census period, we calculated the proportion of colonies surviving to the next census period. This calculation yielded 10 between-census estimates of survival for each of the four treatment groups. To account for potential seasonal differences in mortality (typically winter-biased for this species in natural populations; Herbers and Johnson 2007; Mitrus 2013), we took the ratio of the ‘away’ treatments to the ‘home’ treatments as our focal response variable. That is, for each census point, we divided the urban-origin transplanted into the rural environment between-census survival by the urban-origin transplanted into the urban environment between-census survival. We did the same for the rural-origin transplanted into the urban environment between-census survival, dividing by the rural-origin transplanted into the rural environment between-census survival at each census point. We then modeled these ratios as a function of season, a two-level factor including summer and winter seasons, a two-level factor of treatment (urban to rural / urban to urban, versus rural to urban / rural to rural), and their interaction. We constructed this model using the *glmmTMB* function with a Gaussian error distribution as the distribution of the response variable and visual inspection of the model residuals indicated this approach was valid.

For our ‘season’ predictor variable, we lumped census points into two categories: summer to early fall, and winter to early spring. The assignment of census points to summer versus winter categories was verified by the average monthly temperatures between each census interval (temperature data were based on NOAA GSOM): summer census intervals had a mean ± 1 SD of 20.3 ± 2.22 °C, and the winter census intervals had a mean ± 1 SD of 8.69 ± 1.68 °C. Owing to strong concordance between temperature means and extremes (**Fig. 1**), we focused our analyses of seasonal bias in mortality to average monthly temperature.

#### Fecundity

Because the most relevant estimate of fitness via fecundity for an acorn ant colony is the total number of alates (reproductive ants) produced, we summed the total number of alates produced per colony over the course of the transplant experiment. Colonies received a value of 0 if they either failed to produce alates or the colony died over the course of the experiment. We modeled the total number of alates produced using a generalized linear model with a quasi-Poisson error distribution. As the sole predictor, we included the four-level factor of treatment representing each combination of source population and environment.

#### Colony size

Although our focal analyses concerned more comprehensive metrics of fitness including survival and fecundity, we also examined the potential for treatment and census effects on colony size, including the number of workers and number of brood. We took a similar modeling approach as to that used in the cumulative survival model, except that we used the Poisson error distribution and included a random intercept for colony identity as we fit these models as repeated measures at the level of the colony rather than the proportion of colonies that survived at each census point. We performed separate models for workers and brood as well as a model of total colony size, summing numbers of workers and brood.

#### Patterns of phenotypic selection

Although very much a secondary goal of our study, we examined patterns of phenotypic selection on thermal tolerance traits acting in the urban versus rural environments. Note that we pooled data across populations and focused on patterns of selection within a given environment, urban or rural. We used the *gam.gradients* function from the {gsg} library (Morrissey and Sakreida 2014) to estimate directional selection gradients (β) and quadratic selection gradients (γ) for each combination of environment (urban and rural) and thermal tolerance trait (CT_max_ and CT_min_). Separate models were fit for each tolerance trait × environment combination. In each case, the underlying *gam* model was fit using a negative binomial error distribution and a smooth term for the thermal tolerance trait value. Each *gam* model also included a random effect for colony identity since multiple individuals were assessed for thermal tolerance from the same colony. Owing to limited statistical power to estimate selection, we focused our interpretations on the general patterns of phenotypic selection rather than using such analyses for strong inference.

#### Local adaptation meta-analysis

For our effect size, we calculated local adaptation of urban and rural populations relative to the foreign population following the methods of Hereford (2009). To calculate local adaptation, we used the relative fitness of the native population at their home site/treatment minus the relative fitness of the foreign population at the same site/treatment.

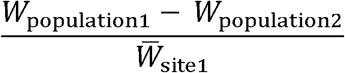

Here *W* represents the mean fitness of each population at site 1 and 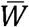 represents the mean fitness of all populations at site 1. This metric is an estimate of selection against migrants into a population with positive values indicating selection against migrants (*i.e.,* local adaptation), and negative values indicating that migrants would have greater fitness than the native population (*i.e.*, maladaptation of the native population). We also calculated the pooled standard errors of the estimates of relative fitness for each population.

We fit mixed-effect meta-analytic models using the *rma.mv* function in the {metafor} library (Viechtbauer 2010) to evaluate (1) evidence for local adaptation to urban and rural habitats (*i.e.* that the metric of local adaptation was > 1); (2) if the magnitude of urban adaptation differed from that of rural adaptation; (3) if local adaptation differed between lab and field experiments; and (4) if local adaptation differed among fitness metrics. Each model was fit separately with a single predictor variable, the squared pooled standard errors (*i.e.,* the measurement error), and species identity as a random effect (which largely overlaps with study identity).

## Results

### Cumulative and census-specific survival across treatments

The model of cumulative survival across the 11 census points and the four treatment groups, including urban-origin population colonies transplanted to the urban environment, rural-origin to the rural environment, urban-origin to the rural environment and rural-origin to the urban environment, revealed a significant effect of treatment on cumulative survival that also depended on the specific census point (treatment main effect: χ^2^ = 28.0, *P* < 0.0001; census point main effect: χ^2^ = 16.9, *P* < 0.0001; treatment × census interaction: χ^2^ = 18.3, *P* = 0.000391) (**Fig. 2**). Post-hoc analyses allowed us to assess the census points at which the trade-off between home treatments (urban-origin to urban environment and rural-origin to rural environment) versus away treatments (urban-origin to rural environment and rural-origin to urban environment) became apparent (**Fig. 1**; **Tables S4,S5**). By the midpoint of the experiment at census 6 (July, 2018), several patterns emerged: higher survival of the urban-origin ants than rural-origin ants in the urban environment; higher survival of the rural-origin ants than urban-origin ants in the rural environment; higher survival of the urban-origin ants in the urban environment compared with the rural environment; and higher survival of the rural-origin ants in the rural environment compared with the urban environment. All such comparisons were statistically significant at census 6 except the urban-origin ants in the rural versus urban environment. It was not until census 8 that the higher survival of the urban-origin ants in the urban versus rural environment was statistically significantly different.

**Figure 2.**
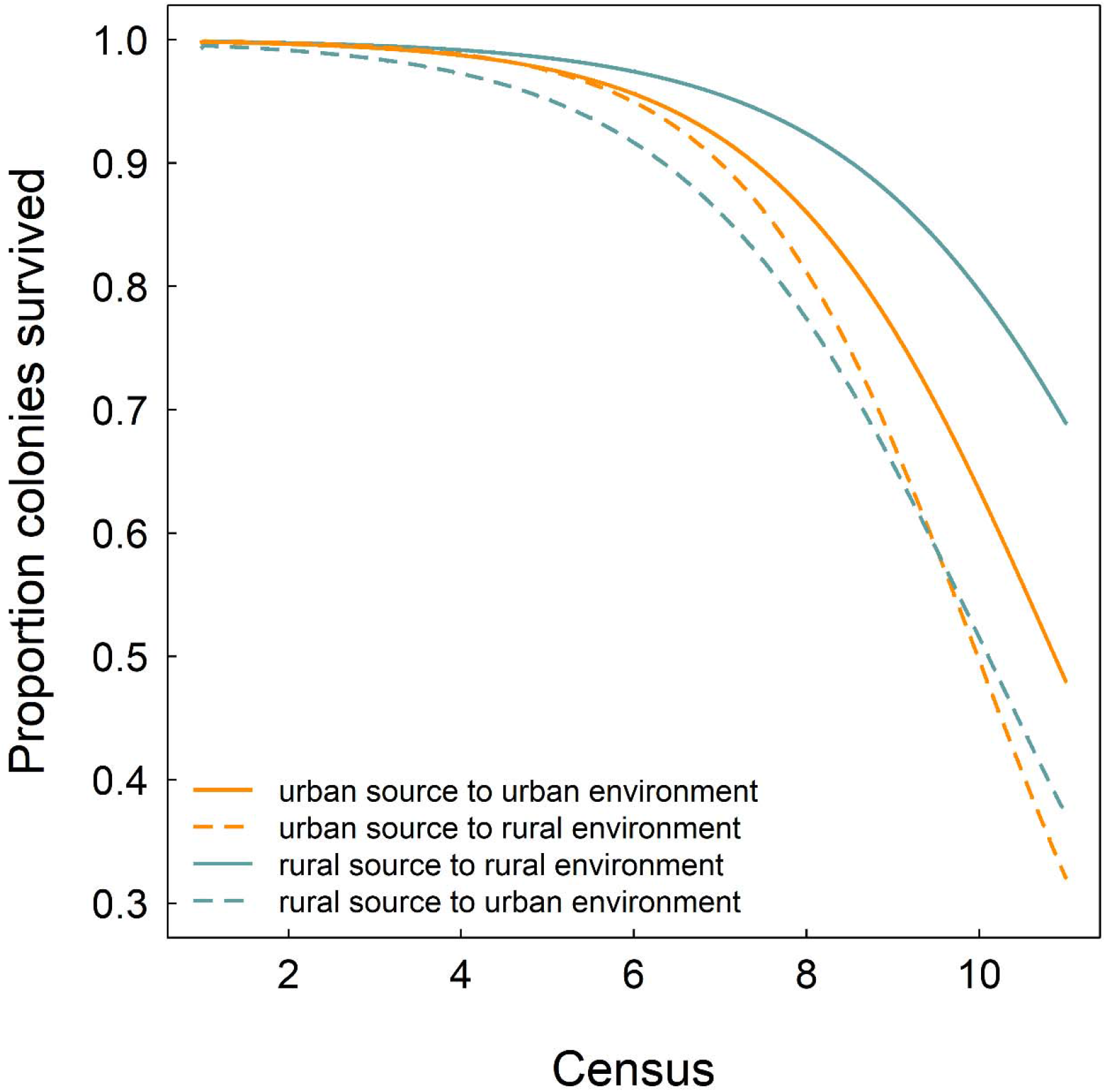
Survival curves (cumulative) over the entire experiment for each combination of source population and environment. Predicted values from a generalized linear model that accounts for temporal autocorrelation.

### Seasonal bias in mortality

Our model of between-census survival revealed a significant interaction between relative survival of away versus home treatments and summer versus winter seasons (season main effect: χ^2^ = 5.29, *P* < 0.0001; population main effect: χ^2^ = 12.8, *P* = 0.000344; season × population interaction:χ^2^ = 39.6, *P* < 0.0001), which prompted us to use post-hoc tests to examine the nature of this interaction effect. We found that, for the rural-origin population colonies, relative survival of away versus home treatments was significantly lower in summer compared with winter (contrast of summer – winter: estimate χ^2^ = −0.0327, SE = 0.0142, *df* = 15, *t* = −2.30, *P* = 0.0362; **Fig. 3**). By contrast, for the urban-origin population colonies, we found that the relative survival of away versus home treatments was significantly lower in winter compared with summer (contrast of summer – winter: estimate = 0.0911, SE = 0.0136, *df* = 15, *t* = 6.71, *P* < 0.0001; **Fig. 3**).

**Figure 3.**
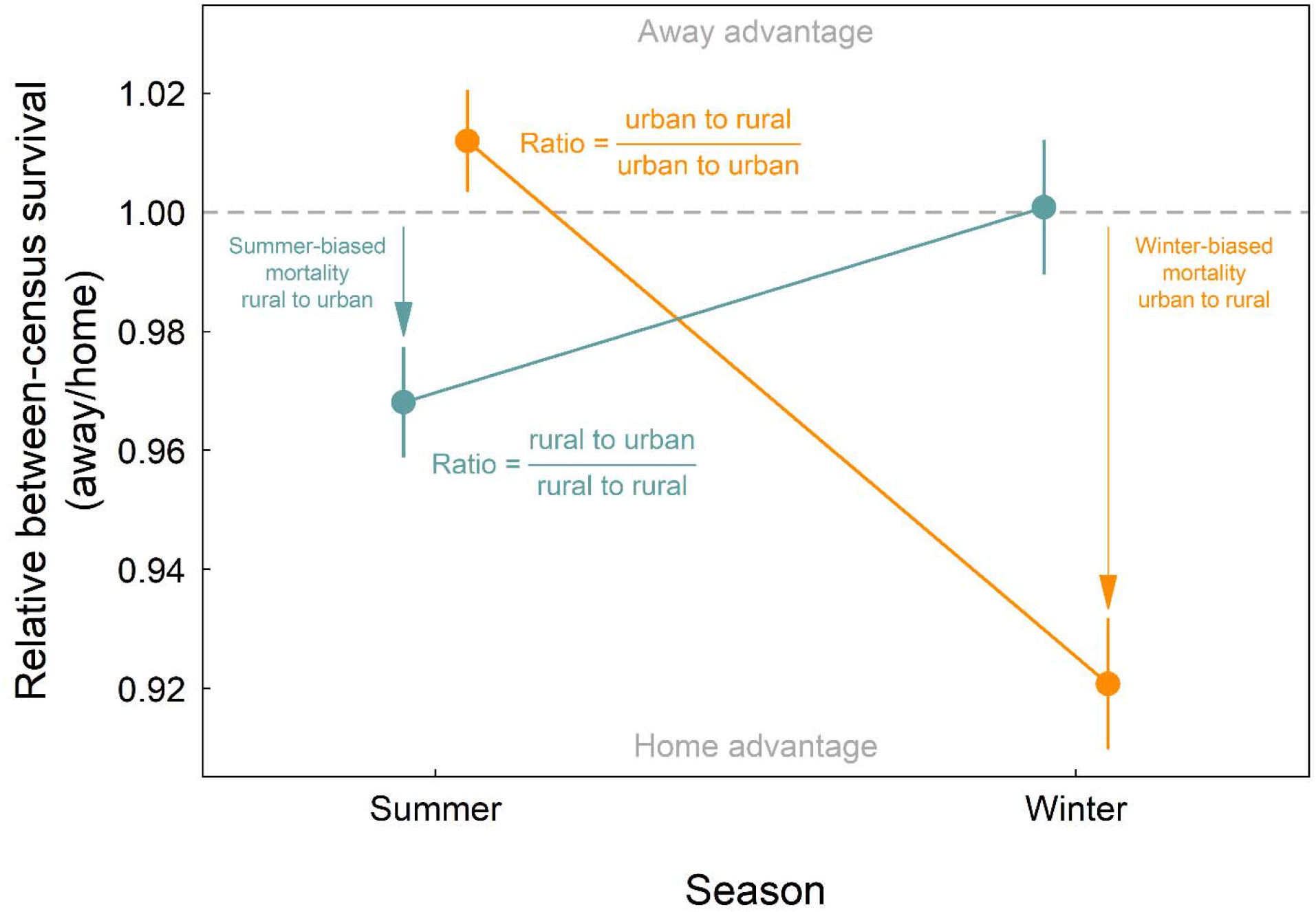
Bias in the seasonal timing of colony survival. Ratio ± 1 SE of away/home treatment between-census survival as a function of the summer versus winter season. Results are presented separately for rural-origin and urban-origin treatments. The dashed gray line indicates equal survival between away and home treatments; below the line indicates a survival advantage of the home treatment, and above the line indicates a survival advantage of the away treatment.

### Fecundity

In some ways, the results for fecundity paralleled the results for survival. Specifically, the rural-origin ants produced more alates than the urban-origin ants in the rural environment (χ^2^ = 12.1, *P* = 0.000497), and the urban-origin ants produced more alates (reproductive ants) in the urban environment than they did in rural environment (χ^2^ = 6.00, *P* = 0.0143) (**Fig. 4**). These results were driven by the complete lack of alate production of the urban-origin ants transplanted to the rural environment. By contrast, we found no evidence for a trade-off with the rural-origin ants transplanted to the urban environment with respect to alate production. We detected no significant difference between the urban-origin and rural-origin ants transplanted to the urban environment (χ^2^ = 1.19, *P* = 0.275); and we detected no significant difference between the rural-origin ants across the urban and rural environments (χ^2^ = 0.580, *P* = 0.447) (**Fig. 4**). The results for fecundity were qualitatively similar between models where all colonies were included versus where only the fecundity of the surviving colonies was considered.

**Figure 4.**
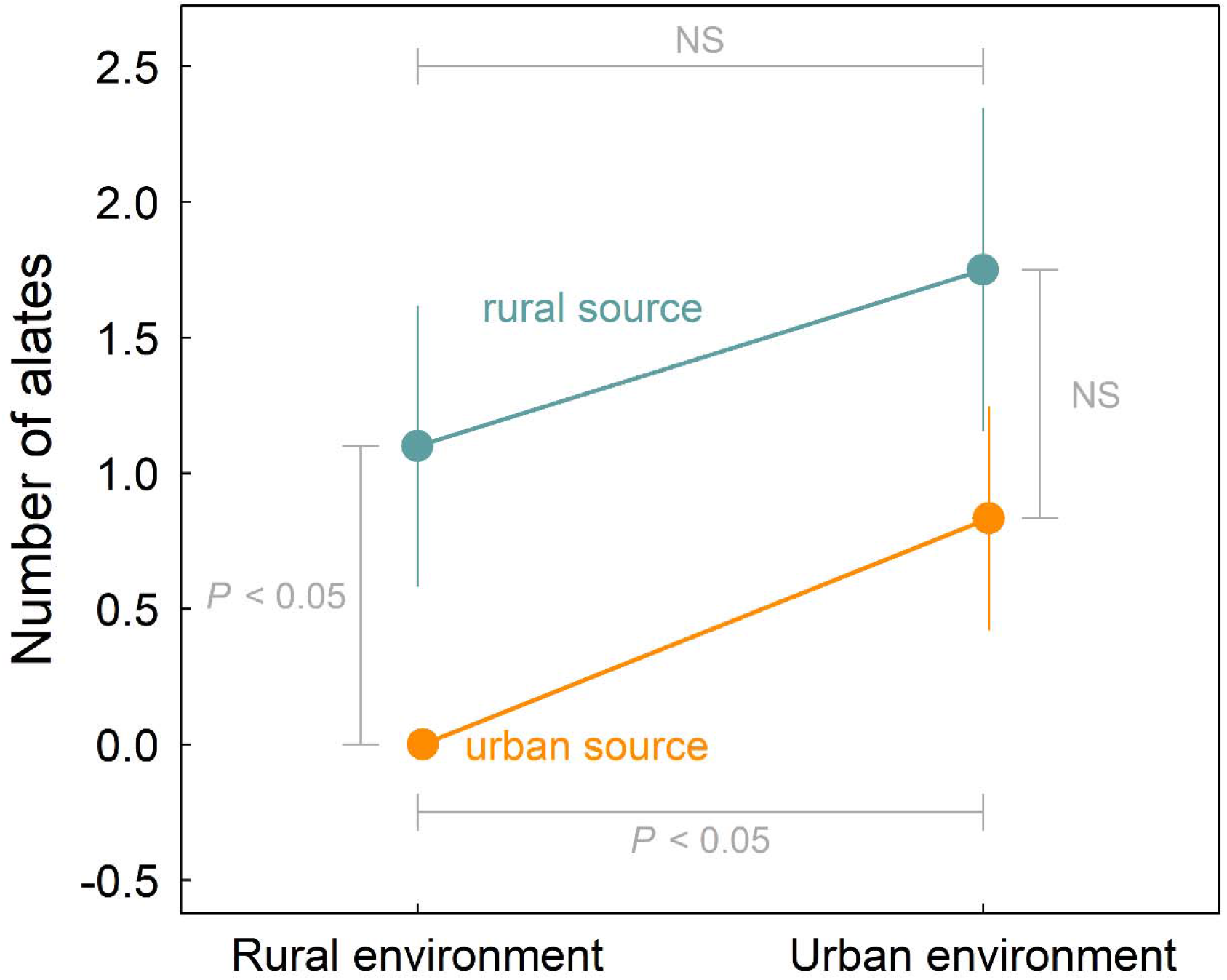
Estimated number of alates (reproductive ants) ± 1 SE produced per colony over the course of the transplant experiment for each combination of source population and environment. Statistical significance of each pairwise contrast of the two source populations within a single environment and of the two environments for a single source population are indicated with brackets and values (NS = non-significant at the *P* < 0.05 level).

### Colony size

Our models of colony size revealed significant variation in the number of brood and workers across census points, with greater colony size in warmer summer months and reduced colony size in cooler winter months (**Figs. S1,S2**). However, we detected no significant effects of treatment (a four-level factor with each combination of source population and environment) on colony size. For simplicity, we present the results of the summed worker and brood numbers: treatment main effect: χ^2^ = 0.0778, *P* = 0.0001; census point main effect: χ^2^ = 8.12, *P* = 0.00437; treatment × census interaction: χ^2^ = 3.27, *P* = 0.353. Individual models of worker number and brood number yielded qualitatively similar results.

### Phenotypic selection on thermal tolerance

Selection gradient analyses revealed a trend of positive directional selection for higher CT_max_ in the urban environment (β ± 1 SE = 0.106 ± 0.156), but a trend of negative directional selection against higher CT_max_ in the rural environment (β ± 1 SE = −0.144 ± 0.245). Trends in selection on CT_min_ were much weaker: we found a weak trend of selection for increased cold tolerance (higher fitness at lower CT_min_ values) in the rural environment and against increased cold tolerance (higher fitness at higher CT_min_ values) in the urban environment (**Table S6**). We focused our interpretations on the directional selection components rather than the quadratic selection components because visual inspection of the jointly estimated directional and quadratic selection coefficients indicated no true fitness minimum or maximum.

### Meta-analysis of local adaptation

Our literature search returned seven studies that met our criteria (**Table 1**). Along with the fitness measures from our current study, this resulted in 27 estimates of local adaptation from six species (14 measured in urban environments and 13 in rural environments). Of these estimates, 64% were consistent with urban local adaption, and 85% for rural local adaptation (*i.e.*, positive values). Across all studies the strength of local adaptation was 0.274 ± 0.142 (*z* = 1.927, *P* = 0.054). The magnitude of local adaptation was greater for rural populations than for urban populations (Q_M_ = 523.706, df = 1, *P* < 0.001) (**Fig. 5)**. Furthermore, the magnitude of rural local adaptation was significantly greater than zero, with rural populations obtaining 34% greater fitness than urban populations in rural environments (mean ± SE: 0.340 ± 0.141; *z* = 2.417, *P* = 0.016). Although it was not significantly different from zero, the magnitude of urban local adaptation also trended positive, with urban populations obtaining 24% greater fitness than rural populations in urban environments (0.241 ± 0.141; *z* = 1.710, *P* = 0.087). However, there was no difference in the magnitude of local adaptation between field and lab studies (Q_M_ = 0.450, df = 1, *P* = 0.502) (**Fig. 5**) or among fitness metrics (Q_M_ = 1.710, df = 2, *P* = 0.425) (**Fig. S3**).

**Figure 5.**
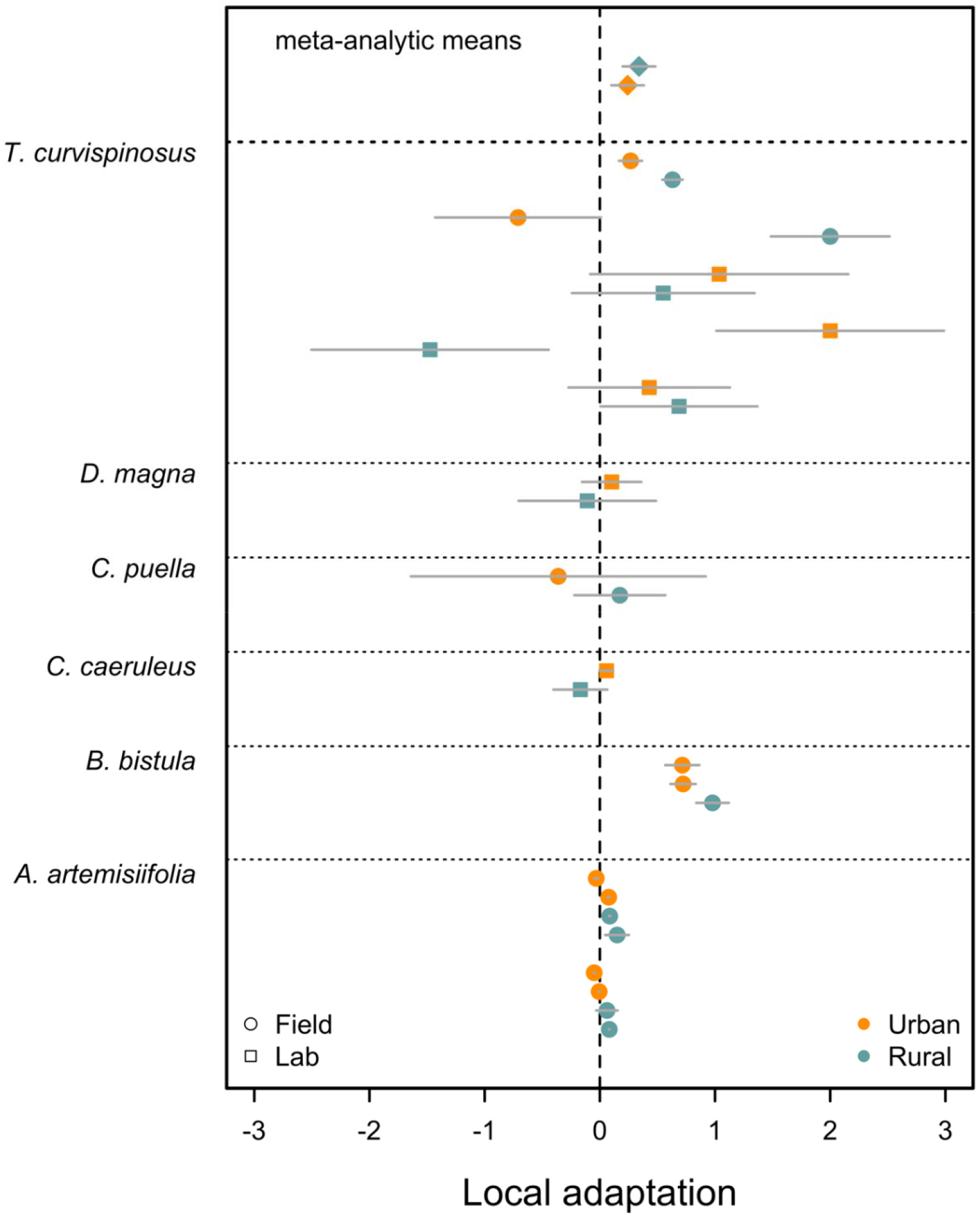
Estimates of local adaptation ± 1 SE from field and lab experiments measuring the fitness of urban and rural populations in urban and rural habitats and manipulated laboratory conditions. Results are grouped by species with the overall meta-analytic means across all studies up top. Positive values indicate greater relative fitness of the native population in comparison to the foreign population (*i.e.*, adaptation) and negative values indicate greater fitness of the foreign population in comparison to the native population (*i.e.*, maladaptation).

## Discussion

Urbanization is a major disruptor to the typical environments under which most organisms evolved (Johnson and Munshi-South 2017; Diamond and Martin 2020a). With substantial development of cities over the last century, urbanized environments provide an excellent venue for exploring contemporary adaptive evolution (Rivkin et al. 2019). We performed a multi-year reciprocal transplant experiment with urban and adjacent rural populations of the acorn-dwelling ant *Temnothorax curvispinosus*. Our study revealed evidence of local adaptation via survival: ant colonies had higher survival in their home environments than the away environments. Mechanistically, these fitness differences appeared to reflect seasonal bias in the timing of mortality, with recently evolved losses of cold tolerance making urban-origin colonies more likely to die during the winter season within the rural environment. In addition, the relatively low heat tolerances in the ancestral, rural population made rural-origin colonies more likely to suffer mortality events during the summer season within the urban environment. Our results showing positive evidence of urban adaptation in acorn ants were further supported by our meta-analysis of fitness differences across urban-rural population pairs. Although there was considerable variation among studies, the overall signal was one of positive support for local adaptation, though the strength of adaptation was greater for the rural compared with the urban populations. Together, our findings provide positive support for contemporary adaptive evolution to cities.

Urban heat island effects are often a key component of urbanization-driven changes to the environment (Diamond and Martin 2020b; Szulkin et al. 2020a). Contemporary evolution of physiological traits to cope with elevated temperatures in cities appears to be fairly repeatable across different taxa (Diamond and Martin in revision, 2020b). Although the evolution of higher heat tolerance, for example, would appear to be adaptive, few studies have shown fitness benefits of the urban population in its home environment compared with the away, rural environment, nor linked the trait with fitness via selection analyses. In water fleas, greater fitness of urban populations compared to rural populations reared in the laboratory suggest that the evolutionary divergence of heat tolerance between these populations is adaptive (Brans and De Meester 2018). Similarly, previous work in the acorn ant system shows evidence of fitness trade-offs in fecundity along with evolutionary divergence in thermal tolerance traits. Phenotypic selection analyses on heat tolerance, for example, via fecundity show positive directional selection for increased heat tolerance in the urban environment (Diamond et al. 2018a). However, these data also come from lab-reared colonies. Field-based evidence of local adaptation to cities generally comes from plant studies. Gorton and colleagues (2018) performed a reciprocal transplant experiment of ragweed plants between multiple urban and rural sites. Although the rural plants generally had higher lifetime reproductive success across rural and urban sites, evidence of phenotypic selection on plants in the urban environment but not the rural environment were suggestive of adaptive evolution. Similarly, Lambrecht and colleagues (2016) found widespread evidence of selection on many hawksbeard plant traits including physiological tolerance via a one-way reciprocal transplant of rural populations into the city. However, this study was not expressly designed to test for local adaptation (rather for patterns of phenotypic selection), as the reciprocal transplant in the other direction (urban into rural) was not performed.

Because acorn ants live within a contained environment and exhibit very low dispersal (Herbers 1990; Stuble et al. 2013), they can also be incorporated into reciprocal transplant studies in a manner analogous to the plant studies described above. This is the approach we took here. By the midpoint of our multi-year reciprocal transplant study, fitness trade-offs via survival started to become apparent (**Fig. 2**). Importantly, these differences manifested in the directions consistent with local adaptation: urban colonies exhibited higher survival in the urban environment compared with the rural environment (and vice versa for the rural colonies), and urban colony survival was higher than rural colony survival within the urban environment whereas rural colony survival was higher than urban colony survival within the rural environment (**Fig. 1**). In effect, survival was greater in the ‘home’ comparisons versus the ‘away’ comparisons. Although we had limited statistical power, the patterns of phenotypic selection via survival on heat and cold tolerance traits suggested that these traits were an important aspect of the adaptation to urbanization in acorn ants. Consistent with the patterns of selection via fecundity from a laboratory study, we found trends of selection for higher heat tolerance in the urban environment and weak trends of selection for improved cold tolerance in the rural environment (**Table S6**). These patterns of phenotypic selection match the directions of trait evolution: urban populations have higher heat tolerance and diminished cold tolerance compared with the rural populations.

Uniquely, the reciprocal transplant study allowed us to examine seasonal changes in survival over the course of the experiment, which we were unable to obtain from previous work in this system within the laboratory setting. We found that the particular evolutionary divergence of thermal tolerance traits in the acorn ant system contributed to seasonal bias when mortality occurred. In particular, we found that the urban populations transplanted to the rural environment were especially prone to mortality in winter, which is consistent with the evolved loss of cold tolerance in the urban population. By contrast, rural populations transplanted to the urban environment were especially prone to mortality in the summer, which is consistent with no evolved compensatory gain in heat tolerance of the ancestral, rural population compared with the high heat tolerance of the urban population (**Fig. 3**). Like most ectothermic species, the plastic response to temperature, while present, also appears to be of insufficient magnitude to fully buffer the rural acorn ant colonies when transplanted into the urban environment (Gunderson and Stillman 2015; Sørensen et al. 2016). The importance of seasonal timing of measuring fitness differences and selection is also reflected in the urban white clover system. In white clover, positive directional selection via biomass (a fitness proxy) for increased cyanogenesis in both urbanized and rural sites was found, despite evolved decreases in cyanogenesis in urban populations (Thompson et al. 2016). Low cyanogenesis is likely due to increased costs associated with cyanogenesis in cold environments characteristic of many cities with urban snow removal which exposes plants to cooler air temperatures than they would experience under the insulating effects of snowpack. Importantly, selection was measured during the summer and therefore might not reflect key selective pressures during the winter season. Notably, no selection via seed set was detected in this experiment. However, this particular experiment was conducted to rule out spatial variation in herbivory as an alternative explanation for temperature-driven changes in cyanogenesis of white clover in response to cities, and thus might not be poised to uncover potential local adaptation of clovers in response to urbanization.

Although we previously found fitness trade-offs in the acorn ant system via fecundity measured in the laboratory (Diamond et al. 2018a), we only found partial support for fecundity-based fitness trade-offs in the field reciprocal transplant study. While rural ants produced more offspring (reproductive, non-sterile) than urban ants within the rural environment, we found no significant difference between rural and urban ant colonies within the urban environment. As a consequence, the urban source population ant colonies had comparable fecundity across both urban and rural environments, whereas rural source population ant colonies exhibited a significant increase in fecundity from the urban to the rural environment (**Fig. 4**). It is important to consider these results in context of the natural history of the acorn ant system. Acorn ant colony fecundity was measured as the number of winged, sexual ants (‘alates’) produced per colony over the course of the experiment (Chick et al. 2019). Often, alate production is strongly tied to seasonal variation in temperature. Specifically, colonies typically produce alates in the later summer months, when temperatures are relatively warm (Alloway et al. 1982; Herbers 1990). It is possible that the urban ant colonies transplanted to the rural environment never receive the correct temperature-photoperiod cue to begin the production of alates, as this environment might be too cold to provide such a cue at the expected photoperiod. By contrast, rural colonies transplanted to the urban environment might have received the warm-season cue to produce alates, resulting in relatively high alate production of rural source population colonies within the urban environment. Because we were able to track alate production across multiple seasons in the reciprocal transplant study and the acorn ants were able to respond to seasonal variation in temperature, this might explain why we did not see the complete trade-off in the field (*i.e.*, due to phenological shifts) as we did in the laboratory. Other possibilities include laboratory food supplementation, field microclimatic variation, or different rates of adaptation for survival and fecundity (*i.e.,* survival may contribute more to fitness than fecundity for these populations, *sensu* Moore and Martin 2019). We detected no significant differences in colony size across the treatments, and so colonies having insufficient numbers of workers to leave the ‘ergonomic’ phase of colony growth and enter the ‘reproductive’ phase (Herbers 1990; Hölldobler and Wilson 1990) seems an unlikely explanation for the alate responses.

Considering our results from both the survival and fecundity components of fitness, a picture emerges that overall, rural-origin ants perform better in the urban environment than the urban-origin ants perform in the rural environment. What might explain the greater fitness trade-off associated with adaptation to the urban environment? Common-garden laboratory studies using some of these same populations have found that while evolving greater tolerance to high temperatures, urban acorn-ants in Cleveland have evolved even greater losses in their cold tolerance (Diamond et al. 2018a; Martin et al. 2019). Whether this is due to relaxed selection on cold tolerance in urban environments, correlated evolution of heat and cold tolerance (*e.g.,* through antagonistic pleiotropy, genetic trait correlations), or another mechanism (*e.g.*, mutation accumulation) is not yet known.

Across a diverse range of organisms and urban-rural pairs, our meta-analysis revealed an overall signal of contemporary local adaptation. The magnitude of local adaptation in our meta-analysis was comparable to broad-scale syntheses of local adaptation in other taxa (Greischar and Koskella 2007; Hoeksema and Forde 2008; Leimu and Fischer 2008; Hereford 2009; Fraser et al. 2011; Sanford and Kelly 2011). More importantly, however, within our urban-rural local adaptation meta-analysis, we detected differences in the strength of local adaptation to urban versus rural environments matching the results from our acorn-ant reciprocal transplant. Specifically, the effect was stronger and statistically significant in the ancestral rural environment compared with the novel urban environment (**Fig. 5**), which was on the borderline of being statistically significant. In effect, the urban populations that are specializing on a novel environment that has never been encountered before in the course of their evolutionary history are actually getting worse in the ancestral environment faster than they are improving in their own environment. Certainly the rural populations have had more generations to evolve, but this does not explain the considerable loss of adaptation for the urban populations to the ancestral rural environment. Longstanding theory predicts that adaptation to a new environment can cause increasing fitness trade-offs to the ancestral environment (Futuyma and Moreno 1988; Fry 1996; Kassen 2002; Schick et al. 2015), which could explain the generally greater fitness trade-offs for urban populations transplanted to their ancestral rural environment. However, in our study, and in other meta-analyses of adaptation (*e.g*., Hereford 2009; Bono et al. 2017), fitness trade-offs across environments were not ubiquitous, with some urban populations even achieving greater fitness across both environments (**Fig. 5**). One hypothesis predicts that divergence across environmentally homogeneous, rather than environmentally heterogeneous environments are more likely to result in fitness trade-offs because any costs of adaptation (*i.e.*, trade-offs) will not be exposed to selection in homogeneous environments (Kassen 2002; Bono et al. 2017). Moving forward, exploring the features of the adaptive landscape within and across cities might help explain the variation in how urban adaptation progresses in response to differing agents of selection.

Regardless of the specific mechanisms underlying urban adaptation, the substantial variation among studies in the magnitude, and sometimes, the direction of the effect of urban local adaptation (**Fig. 5**) has important implications for responses to ongoing global change. Indeed, the presence of such variation suggests that urban populations are broadly still maladapted (not yet on a fitness peak) to the many, rapid stressors imposed by cities, even if they are currently exhibiting adaptive evolutionary responses to urbanization. Thus while it is encouraging that contemporary urban local adaptation is possible, unfortunately, the results of our study join a broad range of evidence showing that the magnitude and rate of anthropogenic environmental change could exceed the capacity of some species to keep pace (reviewed in Diamond and Martin 2020a; and see Radchuk et al. 2019).

Although researchers generally appreciate the value and importance of measuring fitness differences across environments, particularly to assess the adaptive nature of evolutionary divergence, few tests exist because fitness is notoriously difficult to measure (Hendry et al. 2018). Indeed, the studies that are represented in the meta-analysis we performed are a unique subset of taxa that can be easily manipulated in the lab and field, including birds, invertebrates, and plants. Further, of the studies that did measure urban fitness differences, only the ragweed study of Gorton and colleagues (2018), discussed above, measured total fitness under field conditions (**Table 1**). In the acorn ant study described here, it was especially important to measure both survival and fitness components given the somewhat contrasting results and trade-offs among fitness components documented in other systems, including systems across urbanization gradients (Lucas and French 2012). Our study also demonstrated the importance of seasonal differences in fitness trade-offs as single-season results could give misleading inferences. Long-term monitoring also allowed us to uncover critical insights into the nature of selection on thermal tolerance traits of acorn ants. Finally, while our meta-analysis revealed broad evidence of local adaptation for urban evolution systems, the weak effects for urban adaptation specifically suggests that many populations are still in process of adapting to cities. For the acorn ants in particular, it is unclear how future selection will shape acorn ant physiological tolerances with the current evolved loss of cold tolerance in urban populations and persistence of extreme cold temperature events (Diamond and Martin 2020a). Although long-term reciprocal transplant studies are unlikely to be feasible for a number of urban systems, a focused effort to identify systems amenable to reciprocal transplantation and fine-scale temporal monitoring could help to uncover the nature and frequency at which adaptive contemporary urban evolution occurs.

## Authors’ contributions

R.A.M. and S.E.D. planned and designed the study, analyzed the data, carried out the meta-analysis, and wrote the first draft of the manuscript, L.D.C. helped to design the study, collected all experimental data, and contributed to revisions, M.L.G. helped maintain the experiment, collected experimental data, and contributed to revisions.

## Acknowledgements

The Squire Valleevue and Valley Ridge Farm, the Holden Forests and Gardens, and Case Western Reserve University provided access to field sites. An Oglebay Fund grant provided partial financial support. We thank Stephanie Strickler, Crystal Zhao, Noora Khiraoui, Sammi Fremont, and Claire Burchmore for assistance with field and lab work. We also thank Mike Moore for helpful comments on a previous version of this manuscript.

