## Supplemental Material for "In a nutshell, a reciprocal transplant experiment reveals local adaptation and fitness trade-offs in response to urban evolution in an acorn-dwelling ant"

**Supplementary Material**

*Transplant pen construction*

The transplant pens were constructed of aluminum flashing, 60 cm in height with the bottom 15 cm buried in the soil layer to prevent colony emigration. As an additional preventative measure against emigration, the top 5 cm of the inner portion of the aluminum flashing was coated in Insect-a-Slip (BioQuip). Each pen was 50 cm in diameter, which provided colonies with an area greater than the typical foraging distance of this species (Stuble et al. 2013). Transplant pens were haphazardly arranged across the study site. Randomized or gridded designs were not possible owing to the need to avoid tree roots, fallen branches, and other obstructions.

The area within each transplant pen was checked for the possibility of already-present acorn ant colonies that were not part of the experiment, though none were found. We removed all acorns that were already present in the pens and reseeded each pen with five acorns of comparable size and readiness for colonization by acorn ants, *i.e.* the acorns were at least several months old with enough decomposition to allow excavation by workers. The multiple acorns were necessary as acorn ants are known to move nests throughout the season as the colony grows and their original acorns begin to deteriorate (Alloway et al. 1982; Herbers 1989). Acorns were sliced lengthwise with bypass pruning shears and resealed with plastic-coated garden wire to facilitate checking of the acorn ants at each census point. The acorns seeded within each pen were collected from the local site rather than being transferred between sites. Acorns were replaced over the course of the experiment such that five ready-to-inhabit acorns were always available within the transplant pen. The naturally occurring leaf litter layer within the footprint of the chamber was kept intact. Acorn ant colonies were not provided with food supplementation, but rather, were left to scavenge on the existing detritus within the transplant pens following their construction and any litter/detritus material that fell into the pens over the course of the experiment.

**Table S1**. Collection and transplant site details for each colony. Transplant type listed as “from – to”. Colony size is workers plus brood. Thermal tolerances column refers to whether pre-transplant thermal tolerances were performed.

| Unique colony ID | Site of origin | Origin latitude | Origin longitude | Site transplanted to | Transplant latitude | Transplant longitude | Transplant type | Date collected | Date transplanted | Thermal tolerances | Colony size |
| --- | --- | --- | --- | --- | --- | --- | --- | --- | --- | --- | --- |
| TC_014 | Holden Arboretum | 41.6088 | -81.3127 | University Farm | 41.4934 | -81.4263 | rural-rural | 6/6/2017 | 6/16/2017 | yes | 62 |
| TC_026 | Holden Arboretum | 41.6088 | -81.3127 | University Farm | 41.4934 | -81.4263 | rural-rural | 6/6/2017 | 6/16/2017 | no | 17 |
| TC_090 | Holden Arboretum | 41.6088 | -81.3127 | University Farm | 41.4934 | -81.4263 | rural-rural | 6/6/2017 | 6/16/2017 | no | 68 |
| TC_091 | Holden Arboretum | 41.6088 | -81.3127 | University Farm | 41.4934 | -81.4263 | rural-rural | 6/6/2017 | 6/16/2017 | no | 60 |
| TC_092 | Holden Arboretum | 41.6088 | -81.3127 | University Farm | 41.4934 | -81.4263 | rural-rural | 6/6/2017 | 6/16/2017 | no | 80 |
| TC_052 | University Farm | 41.4984 | -81.4245 | Holden Arboretum | 41.6089 | -81.3132 | rural-rural | 5/30/2017 | 6/11/2017 | yes | 108 |
| TC_056 | University Farm | 41.4984 | -81.4245 | Holden Arboretum | 41.6089 | -81.3132 | rural-rural | 5/30/2017 | 6/11/2017 | yes | 105 |
| TC_058 | University Farm | 41.4984 | -81.4245 | Holden Arboretum | 41.6089 | -81.3132 | rural-rural | 5/30/2017 | 6/11/2017 | yes | 68 |
| TC_046 | University Farm | 41.4986 | -81.4227 | Holden Arboretum | 41.6089 | -81.3132 | rural-rural | 5/30/2017 | 6/11/2017 | no | 48 |
| TC_062 | University Farm | 41.4984 | -81.4245 | Holden Arboretum | 41.6089 | -81.3132 | rural-rural | 5/30/2017 | 6/11/2017 | no | 78 |
| TC_012 | University Farm | 41.4986 | -81.4227 | Metroparks Acacia | 41.5034 | -81.4913 | rural-urban | 7/18/2017 | 7/31/2017 | yes | 51 |
| TC_020 | University Farm | 41.4986 | -81.4227 | Metroparks Acacia | 41.5034 | -81.4913 | rural-urban | 7/18/2017 | 7/31/2017 | yes | 120 |
| TC_053 | Holden Arboretum | 41.6088 | -81.3127 | Metroparks Acacia | 41.5034 | -81.4913 | rural-urban | 7/18/2017 | 7/31/2017 | no | 25 |
| TC_064 | Holden Arboretum | 41.6088 | -81.3127 | Metroparks Acacia | 41.5034 | -81.4913 | rural-urban | 7/18/2017 | 7/31/2017 | no | 123 |
| TC_025 | University Farm | 41.4986 | -81.4227 | Botanical garden | 41.5122 | -81.6098 | rural-urban | 5/30/2017 | 6/5/2017 | yes | 100 |
| TC_024 | University Farm | 41.4986 | -81.4227 | Botanical garden | 41.5122 | -81.6098 | rural-urban | 5/30/2017 | 6/5/2017 | no | 46 |
| TC_063 | Holden Arboretum | 41.6088 | -81.3127 | Botanical garden | 41.5122 | -81.6098 | rural-urban | 6/6/2017 | 6/15/2017 | no | 49 |
| TC_023 | University Farm | 41.4986 | -81.4227 | CWRU campus | 41.5042 | -81.6079 | rural-urban | 6/17/2017 | 6/22/2017 | yes | 194 |
| TC_065 | Holden Arboretum | 41.6088 | -81.3127 | CWRU campus | 41.5042 | -81.6079 | rural-urban | 6/17/2017 | 6/22/2017 | yes | 153 |
| TC_018 | University Farm | 41.4986 | -81.4227 | Forest Hill | 41.5197 | -81.5612 | rural-urban | 5/30/2017 | 6/2/2017 | no | 50 |
| TC_049 | Holden Arboretum | 41.6088 | -81.3127 | Forest Hill | 41.5197 | -81.5612 | rural-urban | 6/6/2017 | 6/10/2017 | no | 15 |
| TC_068 | Holden Arboretum | 41.6088 | -81.3127 | Forest Hill | 41.5197 | -81.5612 | rural-urban | 6/6/2017 | 6/10/2017 | no | 36 |
| TC_006 | Ambler Park | 41.4967 | -81.6055 | University Farm | 41.4934 | -81.4263 | urban-rural | 5/22/2017 | 6/6/2017 | no | 116 |
| TC_036 | CWRU campus | 41.5032 | -81.6088 | University Farm | 41.4934 | -81.4263 | urban-rural | 5/22/2017 | 6/6/2017 | yes | 120 |
| TC_035 | CWRU campus | 41.5032 | -81.6088 | University Farm | 41.4934 | -81.4263 | urban-rural | 5/22/2017 | 6/6/2017 | no | 40 |
| TC_041 | Forest Hill | 41.5275 | -81.5737 | University Farm | 41.4934 | -81.4263 | urban-rural | 5/27/2017 | 6/6/2017 | yes | 37 |
| TC_083 | Forest Hill | 41.5275 | -81.5737 | University Farm | 41.4934 | -81.4263 | urban-rural | 5/27/2017 | 6/6/2017 | no | 36 |
| TC_004 | Ambler Park | 41.4967 | -81.6055 | Holden Arboretum | 41.6089 | -81.3132 | urban-rural | 5/22/2017 | 6/11/2017 | yes | 88 |
| TC_011 | CWRU campus | 41.5032 | -81.6088 | Holden Arboretum | 41.6089 | -81.3132 | urban-rural | 5/22/2017 | 6/11/2017 | yes | 114 |
| TC_032 | CWRU campus | 41.5032 | -81.6088 | Holden Arboretum | 41.6089 | -81.3132 | urban-rural | 5/22/2017 | 6/11/2017 | no | 116 |
| TC_081 | Doan Brook | 41.509 | -81.6137 | Holden Arboretum | 41.6089 | -81.3132 | urban-rural | 5/22/2017 | 6/11/2017 | yes | 65 |
| TC_039 | Forest Hill | 41.5275 | -81.5737 | Holden Arboretum | 41.6089 | -81.3132 | urban-rural | 5/27/2017 | 6/11/2017 | no | 35 |
| TC_001 | Ambler Park | 41.4967 | -81.6055 | Metroparks Acacia | 41.5034 | -81.4913 | urban-urban | 7/19/2017 | 7/31/2017 | yes | 78 |
| TC_002 | Ambler Park | 41.4967 | -81.6055 | Metroparks Acacia | 41.5034 | -81.4913 | urban-urban | 7/19/2017 | 7/31/2017 | yes | 110 |
| TC_085 | CWRU campus | 41.5032 | -81.6088 | Metroparks Acacia | 41.5034 | -81.4913 | urban-urban | 7/19/2017 | 7/31/2017 | yes | 83 |
| TC_072 | CWRU campus | 41.5032 | -81.6088 | Metroparks Acacia | 41.5034 | -81.4913 | urban-urban | 7/19/2017 | 7/31/2017 | no | 78 |
| TC_082 | Forest Hill | 41.5275 | -81.5737 | Metroparks Acacia | 41.5034 | -81.4913 | urban-urban | 7/18/2017 | 7/31/2017 | yes | 67 |
| TC_034 | CWRU campus | 41.5032 | -81.6088 | Botanical garden | 41.5122 | -81.6098 | urban-urban | 5/22/2017 | 6/5/2017 | yes | 87 |
| TC_078 | Doan Brook | 41.509 | -81.6137 | Botanical garden | 41.5122 | -81.6098 | urban-urban | 5/22/2017 | 6/5/2017 | yes | 52 |
| TC_074 | Doan Brook | 41.509 | -81.6137 | Botanical garden | 41.5122 | -81.6098 | urban-urban | 5/22/2017 | 6/5/2017 | no | 30 |
| TC_003 | Ambler Park | 41.4967 | -81.6055 | CWRU campus | 41.5032 | -81.6088 | urban-urban | 6/16/2017 | 6/22/2017 | no | 112 |
| TC_040 | Forest Hill | 41.5275 | -81.5737 | CWRU campus | 41.5032 | -81.6088 | urban-urban | 6/17/2017 | 6/22/2017 | yes | 138 |
| TC_077 | Doan Brook | 41.509 | -81.6137 | Forest Hill | 41.5197 | -81.5612 | urban-urban | 5/22/2017 | 5/25/2017 | no | 38 |
| TC_080 | Doan Brook | 41.509 | -81.6137 | Forest Hill | 41.5197 | -81.5612 | urban-urban | 5/22/2017 | 5/25/2017 | no | 51 |

**Table S2**. Seasonal acclimation on thermal tolerance. Effect of Julian day (continuous variable) on thermal tolerance traits from mixed model that accounts for colony-level autocorrelation. The numerator degrees of freedom for the effect of Julian day on thermal tolerance traits is 1 for both urban and rural populations; the denominator degrees of freedom is 13 for urban population and 7 for rural population.

| Source population | Tolerance trait | *F* | *P* |
| --- | --- | --- | --- |
| Urban | CT_min_ | 1.59 | 0.229 |
|  | CT_max_ | 0.650 | 0.434 |
| Rural | CT_min_ | 0.00153 | 0.970 |
|  | CT_max_ | 0.0145 | 0.908 |

**Table S3**. Pre-transplant colony demographic analysis across treatments. In each model, the effect of treatment, a 4-level factor for each combination of source population and transplant environment, on components of colony size is shown. The numerator degrees of freedom for the effect of treatment on initial colony size is 3 and the denominator degrees of freedom is 40.

| Colony size component | *F* | *P* |
| --- | --- | --- |
| Workers plus brood | 0.134 | 0.939 |
| Brood only | 0.800 | 0.502 |
| Workers only | 1.21 | 0.318 |

**Table S4**. Model estimates and standard errors of cumulative survival for each combination of treatment and census point.

| Treatment | Census | Estimate | SE |
| --- | --- | --- | --- |
| Urban-origin to rural environment | 1 | 0.995 | 0.0012 |
|  | 2 | 0.995 | 0.00125 |
|  | 3 | 0.994 | 0.0013 |
|  | 4 | 0.988 | 0.00226 |
|  | 5 | 0.987 | 0.00236 |
|  | 6 | 0.986 | 0.00246 |
|  | 7 | 0.984 | 0.00274 |
|  | 8 | 0.886 | 0.0151 |
|  | 9 | 0.538 | 0.0294 |
|  | 10 | 0.466 | 0.0312 |
|  | 11 | 0.363 | 0.0325 |
| Urban-origin to urban environment | 1 | 0.994 | 0.00144 |
|  | 2 | 0.993 | 0.0014 |
|  | 3 | 0.993 | 0.00137 |
|  | 4 | 0.987 | 0.00223 |
|  | 5 | 0.987 | 0.00219 |
|  | 6 | 0.987 | 0.00214 |
|  | 7 | 0.987 | 0.00221 |
|  | 8 | 0.915 | 0.0119 |
|  | 9 | 0.636 | 0.0267 |
|  | 10 | 0.588 | 0.029 |
|  | 11 | 0.505 | 0.0321 |
| Rural-origin to urban environment | 1 | 0.985 | 0.00282 |
|  | 2 | 0.985 | 0.00273 |
|  | 3 | 0.985 | 0.00268 |
|  | 4 | 0.973 | 0.00522 |
|  | 5 | 0.974 | 0.00488 |
|  | 6 | 0.975 | 0.00454 |
|  | 7 | 0.975 | 0.0045 |
|  | 8 | 0.854 | 0.0188 |
|  | 9 | 0.498 | 0.0287 |
|  | 10 | 0.458 | 0.0302 |
|  | 11 | 0.386 | 0.0311 |
| Rural-origin to rural environment | 1 | 0.995 | 0.00136 |
|  | 2 | 0.995 | 0.00124 |
|  | 3 | 0.995 | 0.00113 |
|  | 4 | 0.992 | 0.00173 |
|  | 5 | 0.992 | 0.00156 |
|  | 6 | 0.993 | 0.00141 |
|  | 7 | 0.993 | 0.00135 |
|  | 8 | 0.954 | 0.00542 |
|  | 9 | 0.781 | 0.0218 |
|  | 10 | 0.754 | 0.0261 |
|  | 11 | 0.698 | 0.0335 |

**Table S5**. Post-hoc comparisons for the cumulative survival model across treatments and census points. Estimates of the contrast (difference), standard errors, *t*-statistics and *P*-values are provided. Results are presented both as population comparisons within an environment type and as environment type comparisons for a given population.

| Type of contrast | Specific contrast | Census | Estimate | SE | *t* | *P* |
| --- | --- | --- | --- | --- | --- | --- |
| Population comparisons within a single environment | Rural-origin to urban environment - urban-origin to urban environment | 1 | -0.00445 | 0.00342 | -1.3 | 0.202 |
|  |  | 2 | -0.00701 | 0.00477 | -1.47 | 0.151 |
|  |  | 3 | -0.011 | 0.00657 | -1.67 | 0.105 |
|  |  | 4 | -0.017 | 0.009 | -1.89 | 0.0678 |
|  |  | 5 | -0.0259 | 0.0123 | -2.11 | 0.0426 |
|  |  | 6 | -0.0388 | 0.0169 | -2.29 | 0.0282 |
|  |  | 7 | -0.056 | 0.0231 | -2.43 | 0.0208 |
|  |  | 8 | -0.077 | 0.0303 | -2.54 | 0.0159 |
|  |  | 9 | -0.0987 | 0.0363 | -2.72 | 0.0104 |
|  |  | 10 | -0.115 | 0.0383 | -3 | 0.00506 |
|  |  | 11 | -0.119 | 0.0388 | -3.07 | 0.00426 |
|  | Urban-origin to rural environment - rural-origin to rural environment | 1 | 7.31E-05 | 0.000666 | 0.11 | 0.913 |
|  |  | 2 | -0.00046 | 0.00103 | -0.444 | 0.66 |
|  |  | 3 | -0.00179 | 0.00178 | -1.01 | 0.321 |
|  |  | 4 | -0.0048 | 0.00323 | -1.49 | 0.146 |
|  |  | 5 | -0.0111 | 0.00593 | -1.88 | 0.0691 |
|  |  | 6 | -0.0237 | 0.011 | -2.16 | 0.038 |
|  |  | 7 | -0.0473 | 0.0203 | -2.33 | 0.0262 |
|  |  | 8 | -0.0881 | 0.0361 | -2.44 | 0.0203 |
|  |  | 9 | -0.151 | 0.0573 | -2.63 | 0.0128 |
|  |  | 10 | -0.232 | 0.0735 | -3.16 | 0.00337 |
|  |  | 11 | -0.314 | 0.0667 | -4.71 | 4.35E-05 |
| Environment contrasts for a single population | Urban-origin to rural environment - urban-origin to urban environment | 1 | 0.000806 | 0.000876 | 0.921 | 0.364 |
|  |  | 2 | 0.001 | 0.00119 | 0.844 | 0.405 |
|  |  | 3 | 0.00103 | 0.00162 | 0.632 | 0.531 |
|  |  | 4 | 0.000502 | 0.00229 | 0.219 | 0.828 |
|  |  | 5 | -0.00139 | 0.00342 | -0.405 | 0.688 |
|  |  | 6 | -0.00622 | 0.0056 | -1.11 | 0.275 |
|  |  | 7 | -0.0167 | 0.01 | -1.67 | 0.104 |
|  |  | 8 | -0.0364 | 0.018 | -2.02 | 0.0513 |
|  |  | 9 | -0.068 | 0.0292 | -2.33 | 0.026 |
|  |  | 10 | -0.108 | 0.038 | -2.84 | 0.00764 |
|  |  | 11 | -0.144 | 0.0403 | -3.57 | 0.00113 |
|  | Rural-origin to urban environment - rural-origin to rural environment | 1 | -0.00518 | 0.00391 | -1.32 | 0.195 |
|  |  | 2 | -0.00847 | 0.00563 | -1.5 | 0.142 |
|  |  | 3 | -0.0138 | 0.00803 | -1.72 | 0.0955 |
|  |  | 4 | -0.0223 | 0.0114 | -1.95 | 0.0595 |
|  |  | 5 | -0.0357 | 0.0163 | -2.19 | 0.0361 |
|  |  | 6 | -0.0563 | 0.0237 | -2.38 | 0.0235 |
|  |  | 7 | -0.0866 | 0.0346 | -2.5 | 0.0175 |
|  |  | 8 | -0.129 | 0.0495 | -2.6 | 0.0139 |
|  |  | 9 | -0.182 | 0.0654 | -2.78 | 0.00899 |
|  |  | 10 | -0.239 | 0.0742 | -3.22 | 0.00286 |
|  |  | 11 | -0.289 | 0.0663 | -4.37 | 0.000118 |

**Table S6**. Selection coefficients (linear, *β*, and quadratic, *γ*) and standard errors.

| Environment | Tolerance trait | Selection gradient | Estimate | SE |
| --- | --- | --- | --- | --- |
| Urban | CT_min_ | *β* | 0.0639 | 0.174 |
|  |  | *γ* | 0.00409 | 0.0549 |
|  | CT_max_ | *β* | 0.106 | 0.156 |
|  |  | *γ* | 0.0113 | 0.0572 |
| Rural | CT_min_ | *β* | -0.0337 | 0.272 |
|  |  | *γ* | 0.00114 | 0.118 |
|  | CT_max_ | *β* | -0.144 | 0.245 |
|  |  | *γ* | 0.0206 | 0.144 |

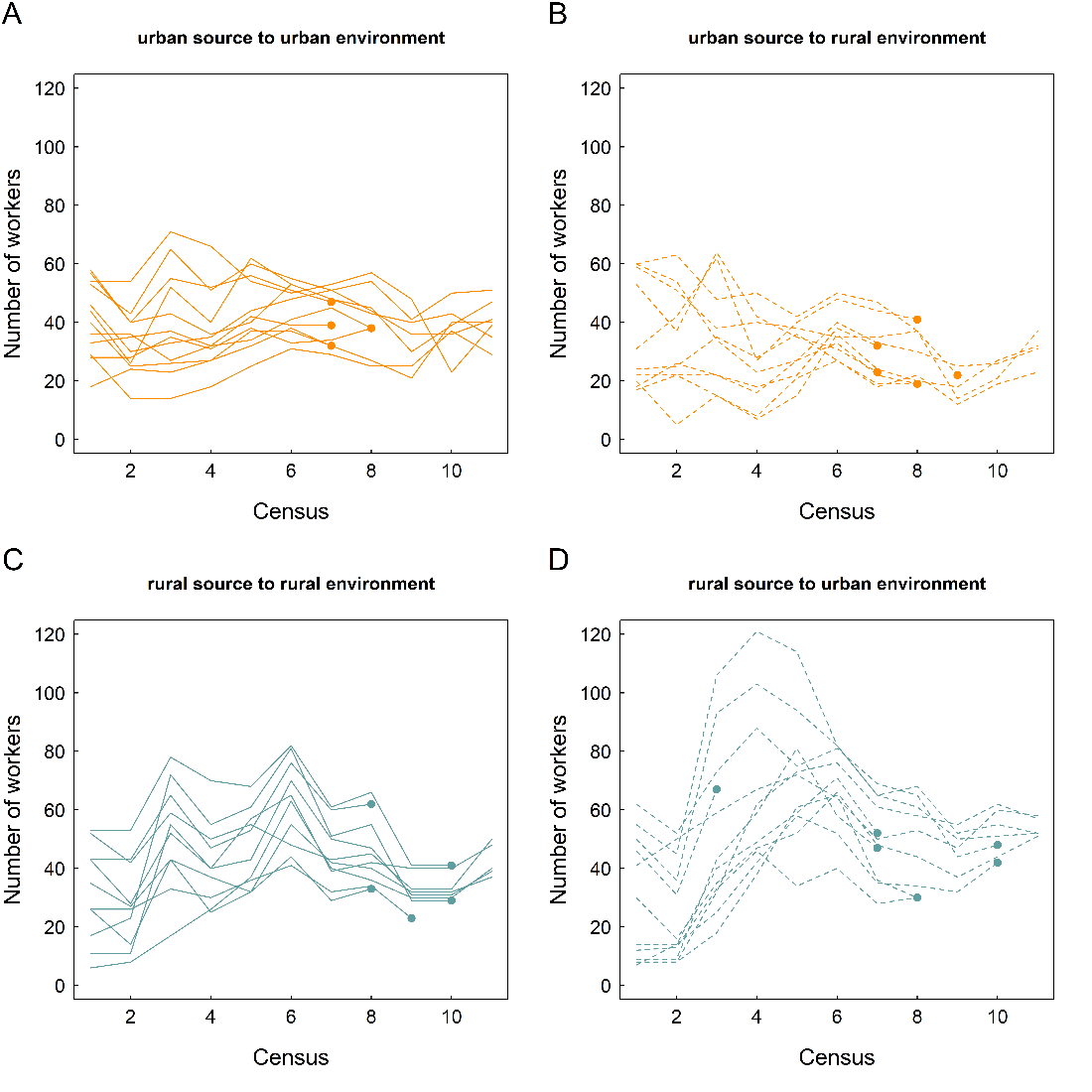

**Figure S1**. Number of workers per colony at each census point. Lines connect responses of individual colonies. Colony mortality events are indicated with points as the last census point at which the colony was alive and able to be counted for workers. (A) urban source to the urban environment; (B) urban source to the rural environment; (C) rural source to the rural environment; and (D) rural source to the urban environment.

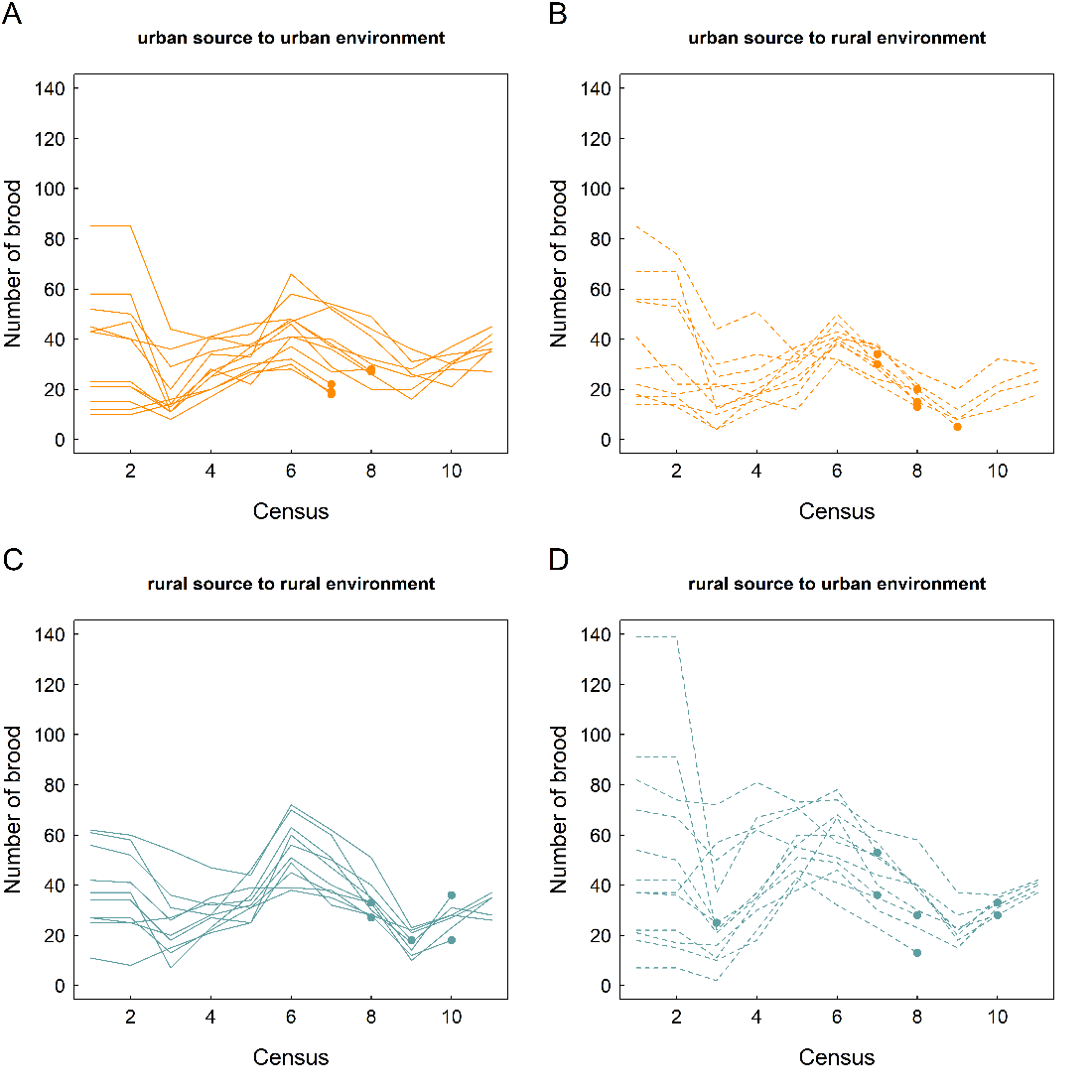

**Figure S2**. Number of brood per colony at each census point. Lines connect responses of individual colonies. Colony mortality events are indicated with points as the last census point at which the colony was alive and able to be counted for brood. (A) urban source to the urban environment; (B) urban source to the rural environment; (C) rural source to the rural environment; and (D) rural source to the urban environment.

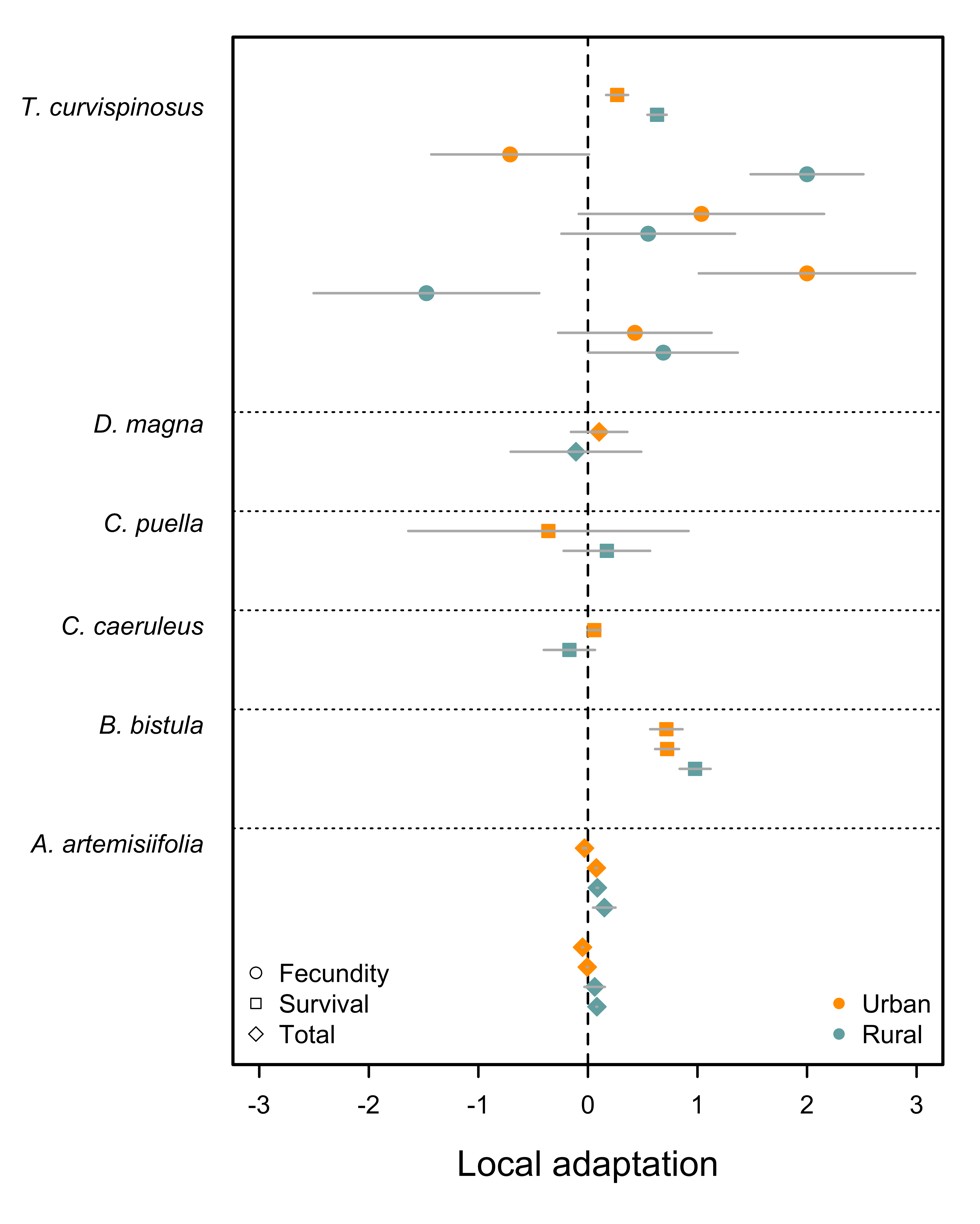

**Figure S3**. Estimates of local adaptation ± 1 SE from field and lab experiments measuring the fitness of urban and rural populations for three fitness metrics. Results are grouped by species. Positive values indicate greater relative fitness of the native population in comparison to the foreign population (*i.e.*, adaptation) and negative values indicate greater fitness of the foreign population in comparison to the native population (*i.e.*, maladaptation).
